# OmniPath: integrated knowledgebase for multi-omics analysis

**DOI:** 10.1101/2025.09.11.675512

**Authors:** Dénes Türei, Jonathan Schaul, Nicolàs Palacio-Escat, Balázs Bohár, Yunfan Bai, Francesco Ceccarelli, Elif Çevrim, Macabe Daley, Melih Darcan, Daniel Dimitrov, Tunca Doğan, Daniel Domingo-Fernández, Aurelien Dugourd, Attila Gábor, Lejla Gul, Benjamin A. Hall, Charles Tapley Hoyt, Olga Ivanova, Michal Klein, Toby Lawrence, Diego Mañanes, Dezső Módos, Sophia Müller-Dott, Márton Ölbei, Christina Schmidt, Bünyamin Şen, Fabian J. Theis, Atabey Ünlü, Erva Ulusoy, Alberto Valdeolivas, Tamás Korcsmáros, Julio Saez-Rodriguez

## Abstract

Analysis and interpretation of omics data largely benefit from the use of prior knowledge. However, this knowledge is fragmented across resources and often is not directly accessible for analytical methods. We developed OmniPath (https://omnipathdb.org/), a database combining diverse molecular knowledge from 168 resources. It covers causal protein-protein, gene regulatory, miRNA, and enzyme-PTM (post-translational modification) interactions, cell-cell communication, protein complexes, and information about the function, localization, structure, and many other aspects of biomolecules. It prioritizes literature curated data, and complements it with predictions and large scale databases. To enable interactive browsing of this large corpus of knowledge, we developed OmniPath Explorer, which also includes a large language model (LLM) agent that has direct access to the database. Python and R/Bioconductor client packages and a Cytoscape plugin create easy access to customized prior knowledge for omics analysis environments, such as scverse. OmniPath can be broadly used for the analysis of bulk, single-cell and spatial multi-omics data, especially for mechanistic and causal modeling.

**Graphical Abstract:** 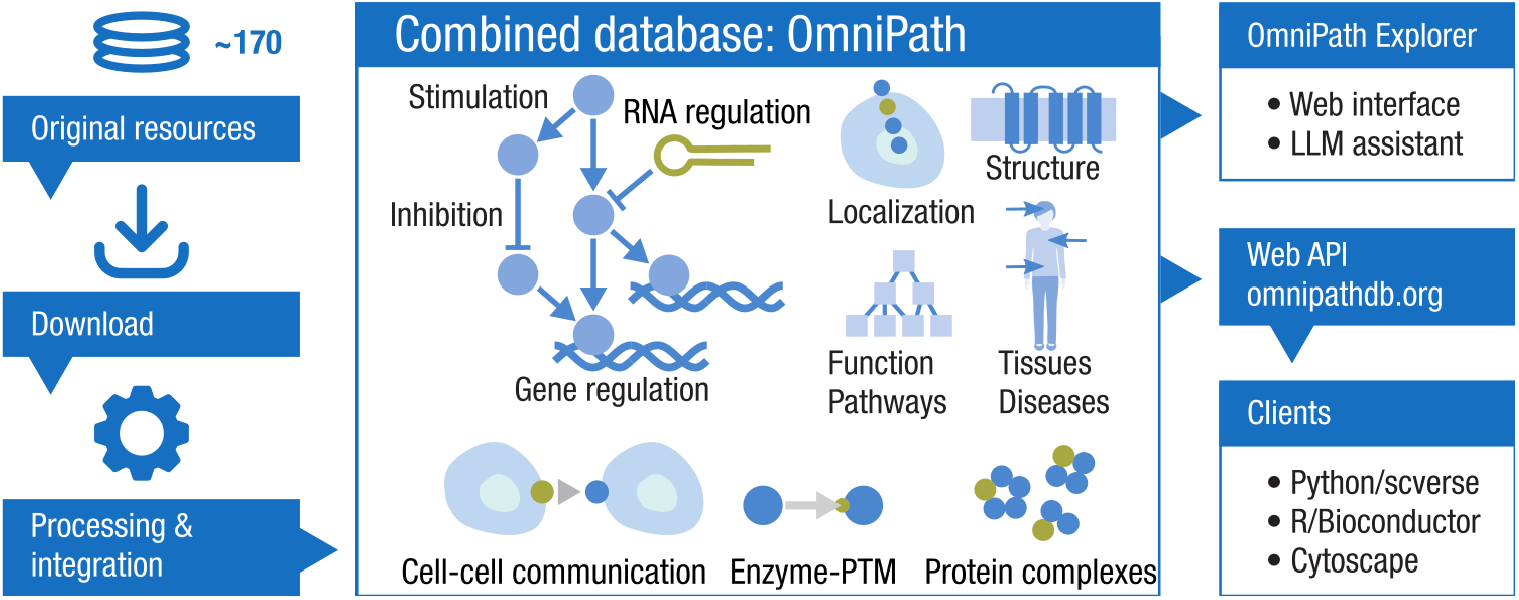

## Introduction

Bulk, single-cell, and spatial omics technologies provide rich information for understanding biological processes, but interpreting molecular mechanisms and their deregulation in disease remains a challenge. The use of prior knowledge within analytical methods largely expands the extractable information and broadens the scope of testable hypotheses. In particular, by estimating the activity(1) of key processes using prior knowledge of signatures—pathway(2), transcription factor (TF; 2, 3), kinase(5), receptor(6) or ligand-receptor(7) activities—the biological interpretability is greatly enhanced and potential causal drivers can be more easily identified, while the dimensionality of the data is reduced, increasing the statistical power(8). Furthermore, the integrated outcome of these estimations can be connected by a variety of network inference methods(9, 10) to derive context-specific mechanisms. Databases of prior knowledge have therefore become essential resources for omics data analysis.

Making prior knowledge available for analysis pipelines is a challenge on its own: it is scattered across many databases and it is not clear which ones are the best suited for the application at hand. Furthermore, each requires different and often significant effort to input into the tools. Excellent original(11–14) and combined(15) databases exist, each with a different focus. Nevertheless these are often limited by various caveats, such as lack of domain knowledge, coverage required by analysis workflows, in particular literature curated, as well as causal interactions, and ease of integration to analysis tools. This challenge prompted us to develop OmniPath, a database combining a growing number of diverse, curated and complementary resources. When first published(16), it only covered causal signaling interactions, and over the years it expanded to include TF-target (regulons) and enzyme-PTM relationships, molecular function and localization, intercellular communication, protein complexes, and various other types of molecular knowledge, as described in a subsequent publication(17). Since then, we included 65 new resources, developed an interactive web page, extended the features of the web API (including provenance details and license support), implemented a higher performance server and added numerous convenient utilities to the Python and R client packages (e.g. identifier and orthology translation). Here we present these novelties and briefly outline our ongoing and future development plans.

## Data content

OmniPath consists of five major database domains: *interactions, enzyme-substrate, complexes, annotations* and *intercellular* (Fig. 1A). These *domains* are integrated databases that we build by combining source databases—what we refer to as *resources* (for a complete list of resources, see Table S1); while within the domains we also define application-specific subsets of resources, which we call *datasets*. Licensing conditions are included for each resource, with the majority of them being available for commercial use (Fig. 1B-C, Table S1). We also classified resources by their maintenance status, i.e. updates happening frequently, infrequently or never. Strikingly, the majority of *interaction* resources and literature references come from databases which have been published only once without subsequent updates (Fig 1B-D, Table S1). In the integrated database multiple resources supporting the same record suggest a higher confidence, while combining the unique content from each resource increases the coverage. In the *interactions* domain only 22 % of the records are unique to a single resource, while in the *complexes* domain more than 60 %, largely due to a lack of agreement between complex prediction methods (Fig 1E).

**Figure 1.**
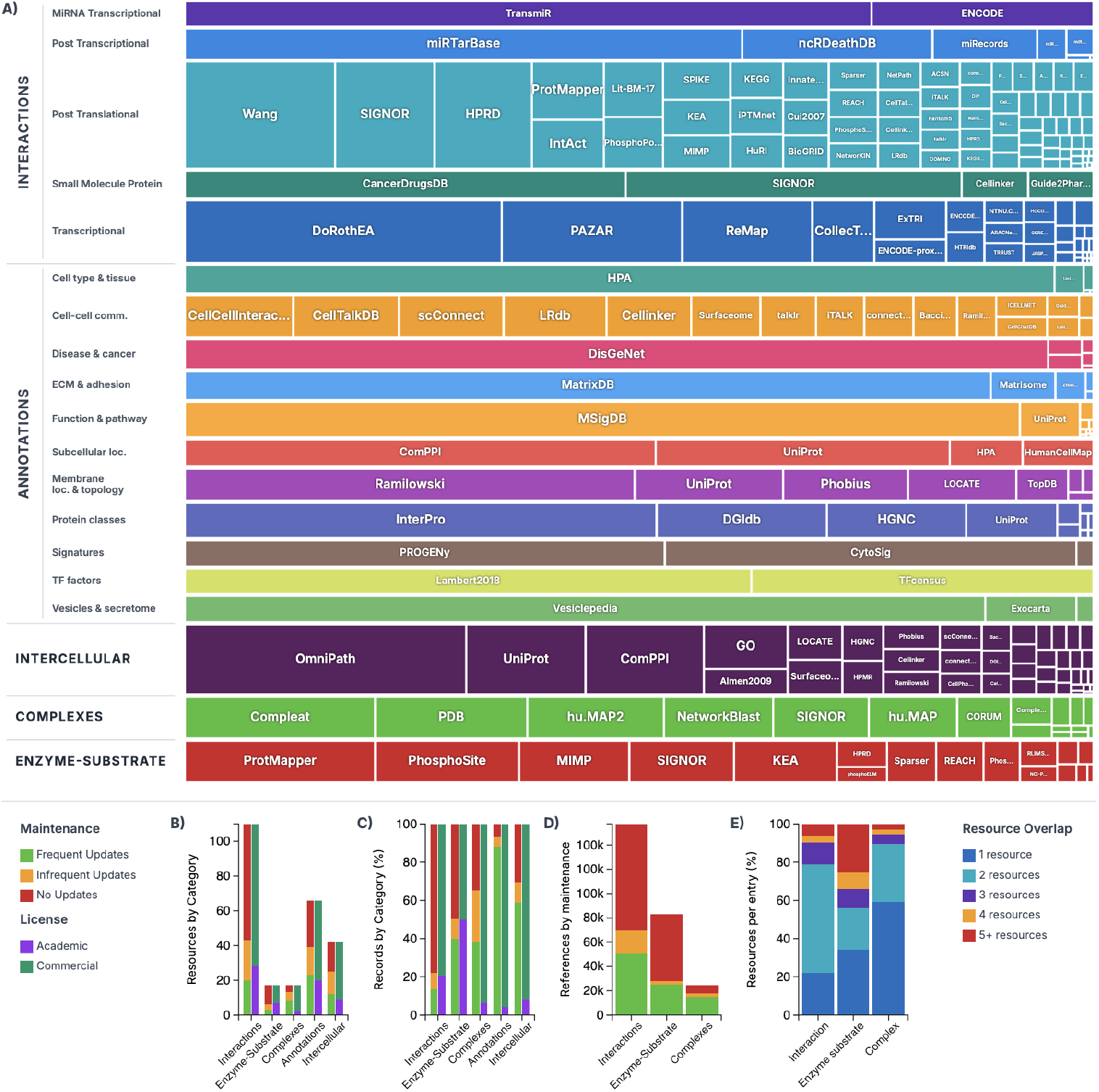
Data content of OmniPath. A) Resource distribution across database categories - Treemap showing relative sizes of individual resources within their database domain and subcategory. The size of each cell is proportional to the number of records. B) Number of resources in each database domain, by maintenance and license categories. C) Percentage of records by database domain and by maintenance and license category. D) Literature reference count by database and maintenance status. E) Resource overlap within the databases: percentage of entries appearing in 1, 2, 3, 4, or 5+ resources. Interactive versions of these visualizations are available at https://explore.omnipathdb.org/.

The *interactions* domain contains 1,419,006 unique molecular interactions from 115 different resources: it covers signaling, gene regulatory (TF-target gene), miRNA-gene, and a limited set of drug-target interactions. Within these interaction types, various datasets are available: the core *omnipath* dataset contains literature-curated, causal PPIs (protein-protein interactions), while further interactions without literature references are provided in separate datasets: *pathwayextra* (causal), *kinaseextra* (kinase-substrate) and *ligrecextra* (ligand-receptor). The largest sources of curated causal signaling interactions are the SIGNOR(11), SignaLink(18) and SPIKE(19) databases (Fig. 1A). Within TF regulons, the *tf_target* dataset contains literature-curated TF-target interactions, while *collectri* and *dorothea* datasets represent comprehensive gene regulatory networks from CollecTRI(3) and DoRothEA(4), respectively, compiled from several curated, high-throughput and predicted sources. Each interaction includes information about its direction and whether it has a stimulatory or inhibitory effect. Hereafter, we refer to this information as “causality”, as they are direct, physical interactions with known biochemical background, though we note that it reflects only putative causal effects.

The *enzyme-substrate* domain is a collection of 115,215 enzyme-PTM interactions from 18 resources. Each record describes the residue, site and type of modification, most predominantly phosphorylation, followed by dephosphorylation and acetylation. PhosphoSitePlus(20) contributes the majority of curated evidence, complemented by further curated and prediction-based resources. The *complexes* database enumerates 52,086 human protein complexes, integrating 18 resources, annotated with stoichiometry and literature references. CORUM(21) and Complex Portal(22), are the primary sources of curated complexes, accompanied by several smaller or prediction-based resources.

The largest database domain in OmniPath is the *annotations* with its 5,895,462 entries, providing a broad variety of protein and gene function, localization, structure, and expression information. This includes pathway memberships, roles in biological processes and diseases, e.g. the functional gene sets from MSigDB(23); various classifications, such as the protein families from HGNC(24); protein localizations—e.g. CSPA(25) to annotate cell surface proteins; and weighted functional signatures, such as the pathway response scores from PROGENy(2) or cytokine responses from CytoSig(26). The data extracted from the 67 resources (Table S1) is provided as it is, without integration across resources.

The *intercell* domain integrates annotation resources into a curated atlas of cell-cell communication. Using the function, localization and structure annotations described above, it classifies proteins into categories such as ligand, receptor, transporter, matrix protein, or secreted enzyme, and tags them with membrane associations and subcellular localization(17). In the R and Python clients, these annotations can be merged with molecular interactions, enabling application-specific customization, for example, by establishing tissue-specific ligand-receptor networks(27).

The *interactions* and *enzyme-substrate* database domains, built with human data, are translated to mouse and rat by orthologous gene pairs. For this translation, we used NCBI HomoloGene(28), Ensembl(29) and the Orthologous Matrix (OMA; 9). Our translation utilities are available in the *pypath* and *OmnipathR* packages, allowing translation to other organisms.

### Web page

With the current update, we introduce *OmniPath Explorer*, an interactive web application (https://explore.omnipathdb.org/) to access the OmniPath resource.

*OmniPath Explorer* consists of three parts: *interactions, annotations*, and *chat* (Fig. 2A-C).

**Figure 2.**
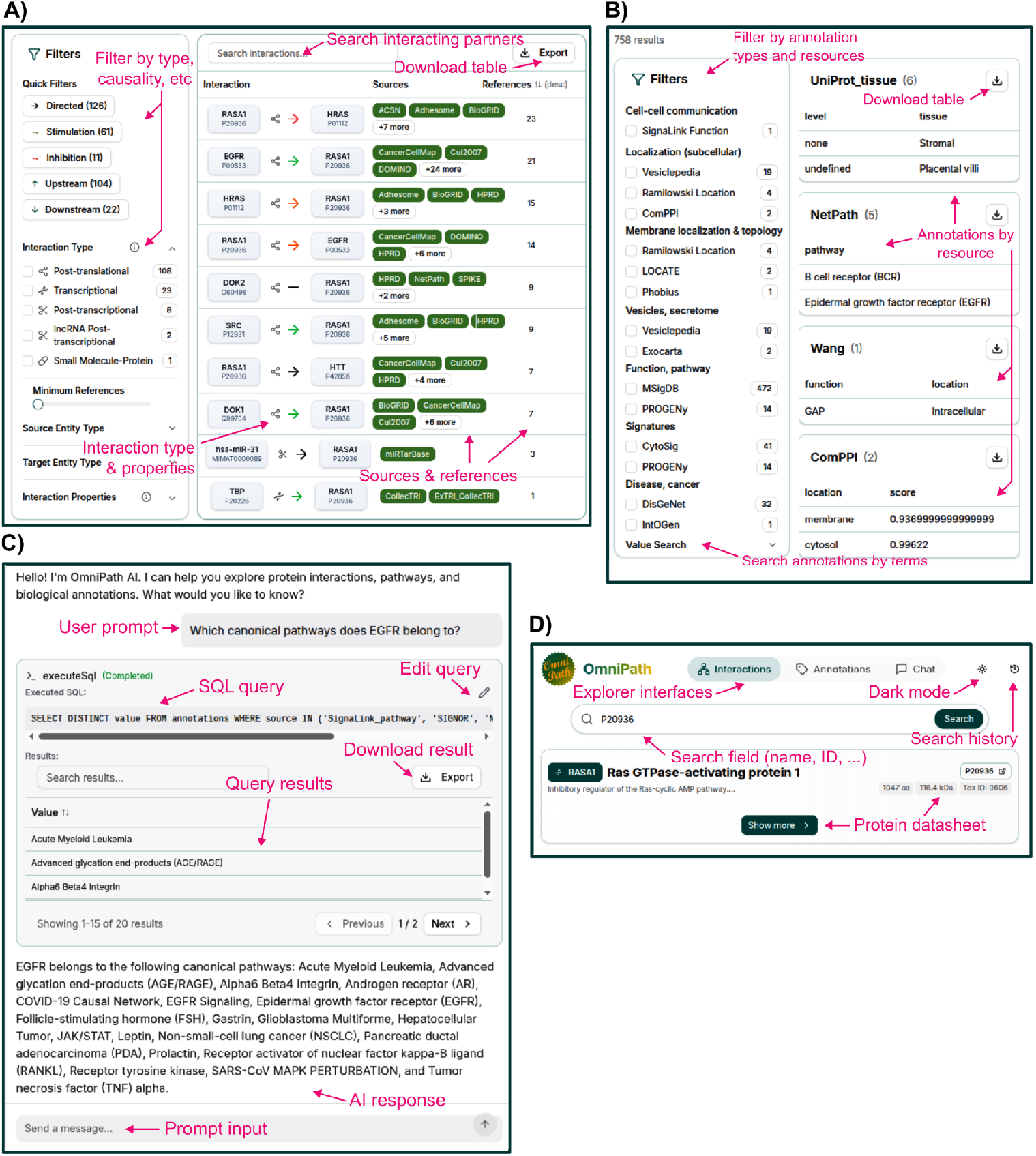
The OmniPath Explorer web application. A) Interaction browser. B) Annotation browser. C) Chat interface to the OmniPath LLM agent. D) Main page search and menu; protein datasheet.

The first two allow users to search for protein and gene names and explore the data through an easy-to-use but comprehensive graphical interface. In the *interactions* part, all partners of a given molecule appear as a list, with the interaction type and causality encoded by colors and symbols. The provenances are also included for each interaction, with links to the original databases and articles in PubMed. The left side control panel enables filtering interactions by type, causality, amount of evidence and other properties (Fig. 2A). The *annotations* part groups the resources by topic, and presents data from the selected ones in tabular format. Alternatively, free text search in annotation records is also available (Fig. 2B). Both in *interactions* and *annotations* view, information about the protein of interest is presented at the top, which includes a basic overview from UniProt(31): the organism, molecular weight, polypeptide length, and literature statements about the functions, with links to PubMed; and in expandable further sections: localization, function, classification, Gene Ontology, PTM information, and links to further resources (Fig. 2D).

The *chat* part integrates an LLM assistant that accepts natural language questions, writes and executes SQL queries, and interprets the results to provide answers. The generated SQL queries and their output are shown alongside the answers, these can be checked for correctness, edited and exported as tables (Fig. 2C). The LLM is provided with an expanding set of query templates which guide it to formulate queries. This enables non-computational users to explore the database content and flexibly access and integrate data across multiple tables or resources.

### Web API

The web service serves data in tabular or JSON format, and consists of five main endpoints (query types), corresponding to the five database domains of OmniPath. It supports the filtering of records by practically any of the variables: by resources, molecules, organisms, and other variables specific for the database domains, e.g. filter TF regulons according to specific confidence levels(4), or PTM residues at the enzyme-substrate domain, or cell-cell communication roles at the *intercell* domain. Optional columns can be selected by the *fields* parameter. *Annotations* are returned as a long-format data frame and require pivoting into wide format, which is supported by the client packages.

We recently added two new columns to the *interaction* records, both containing JSON blobs. The *extra_attrs* column includes resource-specific interaction attributes, such as the mechanism or detection method of the interaction. The client packages are able to extract specific variables from the JSON blob to data frame columns. The *evidences* column contains the provenance information in full detail. This enables the client packages to do precise filtering, e.g. discarding all information that is not licensed for commercial use. License-based filtering is also supported by the web API’s *license* parameter.

Causality of *interactions* is represented by three columns *(is_directed, is_stimulation, is_inhibition)*; in addition, *“consensus”* alternatives of these columns provide a majority vote across all resources.

The web service also features a few auxiliary queries that provide meta-information about the contents. The *databases* and *datasets* queries return the list of resources and datasets in the *interactions* domain, the *queries* query returns the list of valid parameters for each query, while the *resources* query maps the use of resources within the databases, and also includes license information. The *annotations_summary* and *intercell_summary* queries, for each resource, list all variables and all possible values.

## Python, R and Cytoscape clients

The *omnipath* Python package is available in the PyPI repository (https://pypi.org/project/omnipath/), while the *OmnipathR* R package (https://bioconductor.org/packages/OmnipathR) is part of Bioconductor(33). Query types, interaction types and interaction datasets are represented in the Python package by classes, in the R package by functions. All web service query parameters can be provided as arguments to the *“get”* method of the Python classes and similarly to the functions in the R package. The results are returned as data frames. Both packages provide utilities to pivot the annotation data frames from long to wide format, combine networks and annotations, translate data to other organisms by orthologous gene pairs, and update the causality of interactions based on detailed provenance data in the *evidences* column.

Besides the OmniPath client functionalities, *OmnipathR* provides direct access to 26 resources (Table S1), among them several metabolomics related ones, such as MetalinksDB(34), a network of annotated metabolite-protein interactions combining 14 resources, or RaMP-DB(35), a comprehensive resource of metabolite identifiers and structures. *OmnipathR* also comes with a prior knowledge processing toolkit, including translation utilities for 102 identifier types, orthologous gene pair translation (by HomoloGene, Ensembl(29), and OMA(30)), translation ambiguity analysis, handling of organism names, traversal of the Gene Ontology(36) tree, full featured clients to the Ensembl BioMart(29), KEGG(37), and UniProt(32) APIs. It is able to convert *interaction* data frames to igraph(38) networks or deliver them to Bio Model Analyzer(39).

The *OmniPath* Cytoscape app(40) supports the *interactions, enzyme-substrate*, and a limited set of the *annotations* database domains. It imports the data directly into Cytoscape from the datasets and resources specified by the user.

### Implementation

The database build and the web API of OmniPath are implemented in a suite of Python packages (Fig. 3). The *pypath* package is responsible for resource-specific parsing and compiling the combined databases. It features a number of processing utilities, most importantly, the identifier and orthologous gene pair translation. The *pypath*.*inputs* module is a collection of clients for 200 original resources. These clients use the *download-manager* and *cache-manager* packages for robust network transactions and local caching. The *pypath*.*core* module builds the OmniPath databases. All clients and the complete database build are tested daily by an automated pipeline and a status report is published at https://status.omnipathdb.org/. For original resources that became temporarily or permanently inaccessible, we host these on our own server at https://rescued.omnipathdb.org/. Another Python package, the *omnipath-server* loads the databases into PostgreSQL and operates the web service. OmniPath Explorer is a TypeScript application built with the Next.js framework, and uses the same PostgreSQL database as the web API (Fig. 3). The database is updated periodically, with the old versions archived at https://archive.omnipathdb.org/. By default, the LLM agent uses the openly-accessible Google Gemini Flash 2.5 model.

**Figure 3.**
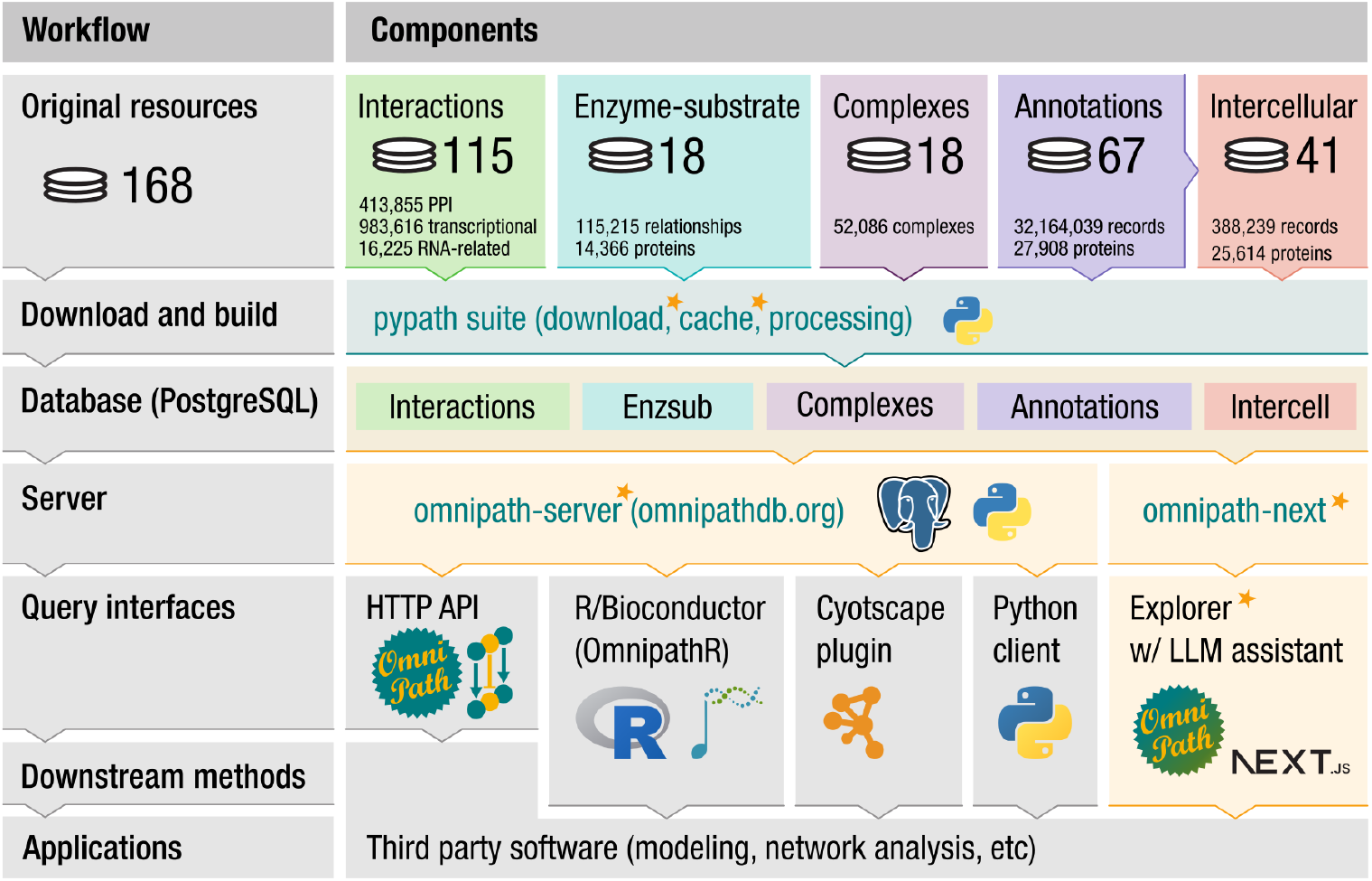
Technical architecture of OmniPath. The combined database, built from 168 different resources by the pypath Python software suite, is publicly available through the HTTP API at https://omnnipathdb.org/. The OmniPath Explorer web app allows interactive browsing with LLM assistance, while client packages for R, Python and Cytoscape provide convenient access and utilities for seamless integration into analysis workflows. The components new or largely renewed in the current update are highlighted in orange boxes and marked with orange stars.

## Discussion

OmniPath is an integrated database combining 168 molecular resources into a single, continuously updated framework, including signed and directed PPIs, enzyme-PTM relationships, ligand-receptor pairs, protein complexes, and extensive functional annotations. By harmonizing data from all these diverse resources, OmniPath facilitates access to prior-knowledge for a broad range of use cases. The OmniPath Explorer presents all evidence for molecular interactions and comprehensive annotations in one place. This allows users to explore interactively the complete knowledgebase and quickly look up specific information and find the most suitable resources for their analysis. The web API and its clients for popular bioinformatics environments (Python, R, Cytoscape) enable effortless creation of customized prior knowledge for diverse applications. With its coverage of small and curated resources, OmniPath also fills a critical gap among other large meta-resources, such as STRING(13) or PathwayCommons(15), and major interaction or pathway databases with original curation effort, like IntAct(12) or Reactome(14).

The integrative design of OmniPath translates into practical impact across diverse tasks in the analysis of omics data. The client packages feature integrations with several downstream analysis tools. For example, integration with the enrichment package Decoupler(1) enables seamless use of signature based transcription factor, pathway, kinase, and cytokine activity estimations in bulk and single cell workflows, including within scverse(41), as well as automated cell type annotations using PanglaoDB(42). Similarly, OmniPath delivers highly customizable ligand-receptor networks directly into the LIANA+(7) cell-cell communication inference framework, also part of the scverse ecosystem. Networks from OmniPath are readily available in the CORNETO(43) network optimization framework, its multi-omics causal variant COSMOS(44) and further network inference methods in NetworkCommons(10), to derive context-specific mechanisms from omics data. For metabolism, OmniPath not only delivers prior knowledge in the MetaProViz R package(45), but also enables connecting knowledge and metabolomics features with its extensive identifier translation utilities. OmniPath is also connected to other prior-knowledge processing systems: the curated knowledge from OmniPath is presented in the INDRA natural language processing system for molecular interactions(46); a script is available to write OmniPath interactions into neo4j importable CSV using the BioCypher library(47); while OmniPath’s prior-knowledge processing toolkit is used in building the CROssBARv2 database(48).

Having a multitude of resources in a uniform format opens the way towards their benchmarking, as it is done, for example, in NetworkCommons(10) to evaluate the performance of network inference methods. Having various interaction types—signaling pathways, transcriptional and miRNA regulation, ligand-receptor, etc—in a uniform network, together with the annotations and the above mentioned inference methods, enables the generation and investigation of complex, multi-layered hypotheses that would be limitedly possible with other resources.

The server-client architecture isolates resource-specific download and processing from analysis workflows, for a reduced complexity and enhanced robustness.

Built-in support for resource license constraints facilitates applications in commercial settings by keeping compliance straightforward. OmniPath Explorer’s LLM agent turns text-based questions into SQL queries. This user-friendly mode of operation assists both new and advanced users to design database queries, while directly answering natural language questions for non-computational users. We consider this to be only a first step towards LLM assistance. Besides expanding the set of query templates, we will work towards a much more structured integration between LLMs and the knowledge contained in OmniPath. We are integrating Omnipath into BioContextAI, a collection of biomedical LLM agents(43), to facilitate integration with LLMs via the model context protocol (MCP).

In our future work, we also plan covering more metabolite, drug, and microbiome-related knowledge, with special focus on causal relationships and functional annotations. We intend to further develop OmniPath Explorer with more views, richer interactivity and information presented in each view. In the Python client side, we have plans to further improve the integration with network inference methods in NetworkCommons and single-cell workflows of the scverse ecosystem.

We remain committed to developing OmniPath as a free open-source resource for the community, particularly for researchers analyzing omics data. We welcome feedback, content suggestions, feature requests, and bug reports, which can be submitted via GitHub (https://github.com/saezlab) or email.

## Data and code availability

The *pypath* Python package is available at https://github.com/saezlab/pypath, the auxiliary packages are linked from this repository. The web API implementation is available at https://github.com/saezlab/omnipath-server, while the *OmniPath Explorer* app at https://github.com/saezlab/omnipath-next. *OmnipathR* is provided at https://github.com/saezlab/OmnipathR, the *omnipath* Python client at https://github.com/saezlab/omnipath and the Cytoscape app at https://github.com/saezlab/Omnipath_Cytoscape. All these components are free software, the database build, web app, server, and Cytoscape app are distributed under GPLv3 license, while *OmnipathR* and *omnipath* client packages under MIT license.

OmniPath data is available at https://omnipathdb.org/ and by the web app at https://explore.omnipathdb.org/, under the licenses of the constituting resources.

Old releases of OmniPath are accessible at https://archive.omnipathdb.org/, and original resources hosted by OmniPath at https://rescued.omnipathdb.org/.

## Supplementary data

Supplementary Data are available at NAR online.

## Acknowledgements

We are grateful to Ömer Kaan Vural, Tennur Kılıç, Ahmet Rifaioglu, and Forrest Hyde for their contributions in the *pypath*.*inputs* module and to Daniele Bottazzi, Jannik Franken, Edwin Carreño, and Youssef Zerta for their ongoing contributions to related software packages.

DT was supported by the Landesinstitut für Bioinformatikinfrastruktur in Baden-Württemberg. JS was supported by the European Union’s Horizon 2020 Programme [Grant no. 965193 (DECIDER)]. NPE acknowledges the support of the German Research Foundation (DFG) through the CRC 1550 “Molecular Circuits of Heart Disease” [Grant No. DFG-508152189]. The work of LG, BB, and TK was supported by the UKRI BBSRC Institute Strategic Programme Food Microbiome and Health [BB/X011054/1 and its constituent project BBS/E/F/000PR13631]. TD, EÇ, MD, BŞ, AÜ and EU were supported by TUBITAK ARDEB 3501 Career Development Program [Project no: 120E531]. SMD was funded by the LiSyM-cancer network supported by the German Federal Ministry of Research, Technology and Space (BMFTR) [031L0257B]. DM acknowledges financial support from Imperial College London through an Imperial College Research Fellowship grant award. CS was funded by SmartCare [03LW0233K].

## Conflict of interests

JSR reports in the last 3 years funding from GSK and Pfizer, and fees and honoraria from Travere Therapeutics, Stadapharm, Astex, Owkin, Pfizer, Grunenthal, Tempus, and Moderna. Work of TK and BB was partially supported by Roche. FJT consults for Immunai Inc., CytoReason Ltd, Cellarity, BioTuring Inc., and Genbio.AI Inc., and has an ownership interest in Dermagnostix GmbH and Cellarity. AV is currently an employee and shareholder of F. Hoffmann-La Roche Ltd.

## References

1. Badia-I-Mompel, P., Vélez Santiago, J., Braunger, J., Geiss, C., Dimitrov, D., Müller-Dott, S., Taus, P., Dugourd, A., Holland, C., Ramirez Flores, R., et al. (2022) decoupleR: ensemble of computational methods to infer biological activities from omics data. Bioinforma. Adv., 2, vbac016.

2. Schubert, M., Klinger, B., Klünemann, M., Sieber, A., Uhlitz, F., Sauer, S., Garnett, M., Blüthgen, N. and Saez-Rodriguez, J. (2018) Perturbation-response genes reveal signaling footprints in cancer gene expression. Nat. Commun., 9, 20.

3. Müller-Dott, S., Tsirvouli, E., Vazquez, M., Ramirez Flores, R., Badia-I-Mompel, P., Fallegger, R., Türei, D., Lægreid, A. and Saez-Rodriguez, J. (2023) Expanding the coverage of regulons from high-confidence prior knowledge for accurate estimation of transcription factor activities. Nucleic Acids Res., 51, 10934–10949.

4. Garcia-Alonso, L., Holland, C., Ibrahim, M., Turei, D. and Saez-Rodriguez, J. (2019) Benchmark and integration of resources for the estimation of human transcription factor activities. Genome Res., 29, 1363–1375.

5. Wirbel, J., Cutillas, P. and Saez-Rodriguez, J. (2018) Phosphoproteomics-Based Profiling of Kinase Activities in Cancer Cells. In Von Stechow, L. (ed), Cancer Systems Biology. Springer New York, New York, NY, Vol. 1711, pp. 103–132.

6. Barsi, S., Varga, E., Dimitrov, D., Saez-Rodriguez, J., Hunyady, L. and Szalai, B. (2025) RIDDEN: Data-driven inference of receptor activity from transcriptomic data. PLOS Comput. Biol., 21, e1013188.

7. Dimitrov, D., Schäfer, P., Farr, E., Rodriguez-Mier, P., Lobentanzer, S., Badia-I-Mompel, P., Dugourd, A., Tanevski, J., Ramirez Flores, R. and Saez-Rodriguez, J. (2024) LIANA+ provides an all-in-one framework for cell-cell communication inference. Nat. Cell Biol., 26, 1613–1622.

8. Schäfer, P.S.L., Dimitrov, D., Villablanca, E.J. and Saez-Rodriguez, J. (2024) Integrating single-cell multi-omics and prior biological knowledge for a functional characterization of the immune system. Nat. Immunol., 25, 405–417.

9. Garrido-Rodriguez, M., Zirngibl, K., Ivanova, O., Lobentanzer, S. and Saez-Rodriguez, J. (2022) Integrating knowledge and omics to decipher mechanisms via large-scale models of signaling networks. Mol. Syst. Biol., 18, e11036.

10. Paton, V., Türei, D., Ivanova, O., Müller-Dott, S., Rodriguez-Mier, P., Venafra, V., Perfetto, L., Garrido-Rodriguez, M. and Saez-Rodriguez, J. (2024) NetworkCommons: bridging data, knowledge and methods to build and evaluate context-specific biological networks. 10.1101/2024.11.22.624823.

11. Lo Surdo, P., Iannuccelli, M., Contino, S., Castagnoli, L., Licata, L., Cesareni, G. and Perfetto, L. (2023) SIGNOR 3.0, the SIGnaling network open resource 3.0: 2022 update. Nucleic Acids Res., 51, D631–D637.

12. del Toro, N., Shrivastava, A., Ragueneau, E., Meldal, B., Combe, C., Barrera, E., Perfetto, L., How, K., Ratan, P., Shirodkar, G., et al. (2022) The IntAct database: efficient access to fine-grained molecular interaction data. Nucleic Acids Res., 50, D648–D653.

13. Szklarczyk, D., Nastou, K., Koutrouli, M., Kirsch, R., Mehryary, F., Hachilif, R., Hu, D., Peluso, M.E., Huang, Q., Fang, T., et al. (2025) The STRING database in 2025: protein networks with directionality of regulation. Nucleic Acids Res., 53, D730–D737.

14. Milacic, M., Beavers, D., Conley, P., Gong, C., Gillespie, M., Griss, J., Haw, R., Jassal, B., Matthews, L., May, B., et al. (2024) The Reactome Pathway Knowledgebase 2024. Nucleic Acids Res., 52, D672–D678.

15. Rodchenkov, I., Babur, O., Luna, A., Aksoy, B.A., Wong, J.V., Fong, D., Franz, M., Siper, M.C., Cheung, M., Wrana, M., et al. (2019) Pathway Commons 2019 Update: integration, analysis and exploration of pathway data. Nucleic Acids Res., 10.1093/nar/gkz946.

16. Türei, D., Korcsmáros, T. and Saez-Rodriguez, J. (2016) OmniPath: guidelines and gateway for literature-curated signaling pathway resources. Nat. Methods, 13, 966–967.

17. Türei, D., Valdeolivas, A., Gul, L., Palacio-Escat, N., Klein, M., Ivanova, O., Ölbei, M., Gábor, A., Theis, F., Módos, D., et al. (2021) Integrated intra- and intercellular signaling knowledge for multicellular omics analysis. Mol. Syst. Biol., 17, e9923.

18. Csabai, L., Fazekas, D., Kadlecsik, T., Szalay-Bekő, M., Bohár, B., Madgwick, M., Módos, D., Ölbei, M., Gul, L., Sudhakar, P., et al. (2022) SignaLink3: a multi-layered resource to uncover tissue-specific signaling networks. Nucleic Acids Res., 50, D701–D709.

19. Paz, A., Brownstein, Z., Ber, Y., Bialik, S., David, E., Sagir, D., Ulitsky, I., Elkon, R., Kimchi, A., Avraham, K.B., et al. (2011) SPIKE: a database of highly curated human signaling pathways. Nucleic Acids Res., 39, D793–D799.

20. Hornbeck, P.V., Zhang, B., Murray, B., Kornhauser, J.M., Latham, V. and Skrzypek, E. (2015) PhosphoSitePlus, 2014: mutations, PTMs and recalibrations. Nucleic Acids Res., 43, D512–D520.

21. Steinkamp, R., Tsitsiridis, G., Brauner, B., Montrone, C., Fobo, G., Frishman, G., Avram, S., Oprea, T.I. and Ruepp, A. (2025) CORUM in 2024: protein complexes as drug targets. Nucleic Acids Res., 53, D651–D657.

22. Balu, S., Huget, S., Medina Reyes, J.J., Ragueneau, E., Panneerselvam, K., Fischer, S.N., Claussen, E.R., Kourtis, S., Combe, C.W., Meldal, B.H.M., et al. (2025) Complex portal 2025: predicted human complexes and enhanced visualisation tools for the comparison of orthologous and paralogous complexes. Nucleic Acids Res., 53, D644–D650.

23. Liberzon, A., Subramanian, A., Pinchback, R., Thorvaldsdóttir, H., Tamayo, P. and Mesirov, J.P. (2011) Molecular signatures database (MSigDB) 3.0. Bioinformatics, 27, 1739–1740.

24. Seal, R.L., Braschi, B., Gray, K., Jones, T.E.M., Tweedie, S., Haim-Vilmovsky, L. and Bruford, E.A. (2023) Genenames.org: the HGNC resources in 2023. Nucleic Acids Res., 51, D1003–D1009.

25. Bausch-Fluck, D., Hofmann, A., Bock, T., Frei, A.P., Cerciello, F., Jacobs, A., Moest, H., Omasits, U., Gundry, R.L., Yoon, C., et al. (2015) A Mass Spectrometric-Derived Cell Surface Protein Atlas. PLOS ONE, 10, e0121314.

26. Jiang, P., Zhang, Y., Ru, B., Yang, Y., Vu, T., Paul, R., Mirza, A., Altan-Bonnet, G., Liu, L., Ruppin, E., et al. (2021) Systematic investigation of cytokine signaling activity at the tissue and single-cell levels. Nat. Methods, 18, 1181–1191.

27. Poletti, M., Treveil, A., Csabai, L., Gul, L., Modos, D., Madgwick, M., Olbei, M., Bohar, B., Valdeolivas, A., Turei, D., et al. (2022) Mapping the epithelial–immune cell interactome upon infection in the gut and the upper airways. Npj Syst. Biol. Appl., 8, 15.

28. Sayers, E.W., Beck, J., Bolton, E.E., Brister, J.R., Chan, J., Connor, R., Feldgarden, M., Fine, A.M., Funk, K., Hoffman, J., et al. (2025) Database resources of the National Center for Biotechnology Information in 2025. Nucleic Acids Res., 53, D20–D29.

29. Dyer, S.C., Austine-Orimoloye, O., Azov, A.G., Barba, M., Barnes, I., Barrera-Enriquez, V.P., Becker, A., Bennett, R., Beracochea, M., Berry, A., et al. (2025) Ensembl 2025. Nucleic Acids Res., 53, D948–D957.

30. Altenhoff, A.M., Train, C.-M., Gilbert, K.J., Mediratta, I., Mendes de Farias, T., Moi, D., Nevers, Y., Radoykova, H.-S., Rossier, V., Warwick Vesztrocy, A., et al. (2021) OMA orthology in 2021: website overhaul, conserved isoforms, ancestral gene order and more. Nucleic Acids Res., 49, D373–D379.

31. The UniProt Consortium, Bateman, A., Martin, M.-J., Orchard, S., Magrane, M., Adesina, A., Ahmad, S., Bowler-Barnett, E.H., Bye-A-Jee, H., Carpentier, D., et al. (2025) UniProt: the Universal Protein Knowledgebase in 2025. Nucleic Acids Res., 53, D609–D617.

32. The UniProt Consortium, Bateman, A., Martin, M.-J., Orchard, S., Magrane, M., Adesina, A., Ahmad, S., Bowler-Barnett, E.H., Bye-A-Jee, H., Carpentier, D., et al. (2025) UniProt: the Universal Protein Knowledgebase in 2025. Nucleic Acids Res., 53, D609–D617.

33. Huber, W., Carey, V.J., Gentleman, R., Anders, S., Carlson, M., Carvalho, B.S., Bravo, H.C., Davis, S., Gatto, L., Girke, T., et al. (2015) Orchestrating high-throughput genomic analysis with Bioconductor. Nat. Methods, 12, 115–121.

34. Farr, E., Dimitrov, D., Schmidt, C., Turei, D., Lobentanzer, S., Dugourd, A. and Saez-Rodriguez, J. (2024) MetalinksDB: a flexible and contextualizable resource of metabolite-protein interactions. Brief. Bioinform., 25.

35. Braisted, J., Patt, A., Tindall, C., Sheils, T., Neyra, J., Spencer, K., Eicher, T. and Mathé, E. (2023) RaMP-DB 2.0: a renovated knowledgebase for deriving biological and chemical insight from metabolites, proteins, and genes. Bioinformatics, 39, btac726.

36. The Gene Ontology Consortium, Aleksander, S.A., Balhoff, J., Carbon, S., Cherry, J.M., Drabkin, H.J., Ebert, D., Feuermann, M., Gaudet, P., Harris, N.L., et al. (2023) The Gene Ontology knowledgebase in 2023. GENETICS, 224, iyad031.

37. Kanehisa, M., Furumichi, M., Sato, Y., Matsuura, Y. and Ishiguro-Watanabe, M. (2025) KEGG: biological systems database as a model of the real world. Nucleic Acids Res., 53, D672–D677.

38. Csardi, G. and Nepusz, T. (2006) The igraph software package for complex network research. InterJournal, Complex Systems, 1695.

39. Benque, D., Bourton, S., Cockerton, C., Cook, B., Fisher, J., Ishtiaq, S., Piterman, N., Taylor, A. and Vardi, M. (2012) Bio Model Analyzer: Visual Tool for Modeling and Analysis of Biological Networks. In Computer Aided Verification (CAV) 2012. LNCS 7358, pp. 686–692. Springer Verlag.

40. Ceccarelli, F., Turei, D., Gabor, A. and Saez-Rodriguez, J. (2020) Bringing data from curated pathway resources to Cytoscape with OmniPath. Bioinformatics, 36, 2632–2633.

41. Virshup, I., Bredikhin, D., Heumos, L., Palla, G., Sturm, G., Gayoso, A., Kats, I., Koutrouli, M., Scverse, C., Berger, B., et al. (2023) The scverse project provides a computational ecosystem for single-cell omics data analysis. Nat. Biotechnol., 41, 604–606.

42. Franzén, O., Gan, L. and Björkegren, J. (2019) PanglaoDB: a web server for exploration of mouse and human single-cell RNA sequencing data. Database J. Biol. Databases Curation, 2019.

43. Rodriguez-Mier, P., Garrido-Rodriguez, M., Gabor, A. and Saez-Rodriguez, J. (2025) Unifying multi-sample network inference from prior knowledge and omics data with CORNETO. Nat. Mach. Intell., 7, 1168–1186.

44. Dugourd, A., Lafrenz, P., Mañanes, D., Fallegger, R., Kroger, A., Turei, D., Shtylla, B. and Saez-Rodriguez, J. (2024) Modeling causal signal propagation in multi-omic factor space with COSMOS. BioRxiv, 10.1101/2024.07.15.603538.

45. Schmidt, C., Turei, D., Prymidis, D., Daley, M., Frezza, C. and Saez-Rodriguez, J. (2025) Integrated metabolomics data analysis to generate mechanistic hypotheses with MetaProViz. bioRxiv, 10.1101/2025.08.18.670781.

46. Bachman, J.A., Gyori, B.M. and Sorger, P.K. (2023) Automated assembly of molecular mechanisms at scale from text mining and curated databases. Mol. Syst. Biol., 19, e11325.

47. Lobentanzer, S., Aloy, P., Baumbach, J., Bohar, B., Carey, V., Charoentong, P., Danhauser, K., Doğan, T., Dreo, J., Dunham, I., et al. (2023) Democratizing knowledge representation with BioCypher. Nat. Biotechnol., 41, 1056–1059.

48. Doğan, T., Atas, H., Joshi, V., Atakan, A., Rifaioglu, A., Nalbat, E., Nightingale, A., Saidi, R., Volynkin, V., Zellner, H., et al. (2021) CROssBAR: comprehensive resource of biomedical relations with knowledge graph representations. Nucleic Acids Res., 49, e96.

49. Kuehl, M., Schaub, D.P., Carli, F., Heumos, L., Fernandez-Zapata, C., Kaiser, N., Schaul, J., Panzer, U., Bonn, S., Lobentanzer, S., et al. (2025) Community-based biomedical context to unlock agentic systems. 10.1101/2025.07.21.665729.

